# Increases in invertebrate abundance and shifts in assemblage composition following Rodent Eradication on Lord Howe Island

**DOI:** 10.1101/2025.10.16.682986

**Authors:** Terence O’Dwyer, Maxim W.D. Adams, Thomas E. White, John Porter, Dean Portelli, Nathan Lo, Nicholas Carlile

## Abstract

Invasive species are one of the major threatening processes impacting biodiversity on islands. In particular, introduced rodents represent one of the most serious threats to island ecosystems, affecting a wide range of native plants, vertebrates and invertebrates. While nearly ubiquitous on human-modified islands, the last four decades have seen the advent of targeted rodent eradications, which have generally resulted in positive impacts for biodiversity. Invertebrates, which are crucial to the functioning of island ecosystems, are known to be negatively impacted by rodents, but their response to rodent removal is less well understood. The largest rodent eradication on an inhabited island was undertaken in 2019 on Australia’s Lord Howe Island, which successfully extirpated black rats (*Rattus rattus*) and house mice (*Mus musculus*) more than a century after their introduction. To examine the impacts of rodents on invertebrates on Lord Howe Island, we collected arboreal and terrestrial species both pre- and post-eradication and identified them to Order. Total invertebrate abundance increased after the eradication of rodents, alongside substantial shifts in assemblage composition, however Ordinal diversity did not change significantly. Orders with large increases in abundance included Isopoda and Blattodea, while the abundance of Coleoptera and Polydesmida did not change. In addition, the abundance of large invertebrates, which are presumably subject to stronger rat predation, rose dramatically following rodent eradication. Our results suggest an ecological rearrangement following the relaxation of predation pressure and augment documented evidence of improved biodiversity outcomes for forest tree species, seabirds and land birds.

## 1. Introduction

Oceanic islands support some of the most biodiverse, yet vulnerable, ecosystems on the planet. Despite contributing less than 7% of terrestrial land area, approximately 75% of recorded species extinctions have occurred on islands (Fernández-Palacios et al., 2021; Sayre et al., 2019). Anthropogenic threats such as habitat clearing, introduction of invasive species, pollution and climate change are anticipated to only intensify in the coming century (Whittaker et al., 2023).

Introduced species are one of the most significant – if not the most significant – drivers of biodiversity loss on islands (Fernández-Palacios et al., 2021; Towns et al., 2006). Rodents in particular are known to readily establish on islands and precipitate declines across a broad swathe of taxa (Angel et al., 2009; Banks & Hughes, 2012; Shiels et al., 2014; Towns et al., 2006). Human-commensal rats (*Rattus* spp.) prey on a wide range of species, including mammals, birds, reptiles, amphibians, invertebrates and plants, which often results in long-term degradation of ecosystem functions (Atkinson, 1985; Auld et al., 2010; Campbell & Atkinson, 2002; Cuthbert & Hilton, 2004; Pender et al., 2013; Shaw et al., 2005; Shiels & Drake, 2011; Smith et al., 2002; Towns et al., 2006). It is estimated that over 90% of the world’s islands now harbour rats or mice (*Mus* spp.), and that these species have often been present for decades (Towns et al., 2006).

In the last 40 years, improvements in biological control methods have made it feasible to completely extirpate invasive species from insular habitats. This has effected a new paradigm in island conservation, centred on invasive-predator eradication (Russell & Broome, 2016). To date, 842 successful rodent eradications have been undertaken on islands, and the biodiversity benefits are typically substantial, though variable across taxa (DIISE, 2019; Holmes et al., 2019; Segal et al., 2021). However, the removal of a top-down regulation of island biota by rodents can have unintended or undesirable consequences due to mesopredator release, new competitive dynamics, or release of invasive prey species (Baker et al., 2020; Bird et al., 2024; Bode et al., 2015; Borrelle et al., 2018; Kurle et al., 2021). Furthermore, since the evidence for the historical adverse effects of rodents on islands is often haphazard or even anecdotal (Towns et al., 2006), eradications can provide valuable opportunities to systematically investigate ecological rebound or trophic cascades, and to discern the consequences of historical rodent incursion.

Lord Howe Island (LHI), a small (< 15 km^2^) volcanic relic *ca*. 600 km east of Australia, is a well-documented example of an island adversely affected by introduced rodents. House mice (*Mus musculus*) probably arrived on LHI in the late 1860s (Hill, 1869), while black rats (*Rattus rattus*) arrived in 1918 following the grounding of the *SS Makambo* (Nicholls, 1953). Both species became widespread across the island, inhabiting built and natural environments. In the century that followed, rats were responsible for the extinction of five endemic bird taxa (Hindwood, 1938), the near-extinction of the Lord Howe phasmid (*Dryococelus australis*) (Priddel et al., 2003), and the declines of many additional species (Lord Howe Island Board, 2007); the impacts of mice upon the fauna are less understood. Rats are also likely to have increased the extinction risk for two endemic palm species through predation of fruits (Auld et al., 2010). However, as robust biodiversity surveys were undertaken only after the arrival of rodents on LHI, the true extent of biodiversity loss remains unknown.

In response to these impacts, a rodent eradication program (REP) was carried out on LHI in 2019 to simultaneously extirpate both black rats and house mice. This was the largest such program ever undertaken on an inhabited island (Harper et al., 2020). A comprehensive success check in 2023 found no evidence of mouse or rat presence on LHI, indicating that eradication was successful (Lord Howe Island Board, 2023). In recognition of the opportunity to study ecosystem recovery, as well as to address public and scientific concerns about the cost effectiveness and merits of rodent eradication (LHIB, 2007), the REP included a multi-faceted biodiversity monitoring program to assess outcomes for island flora and fauna. This included monitoring of invertebrate assemblages because of the important roles these taxa play in ecosystem processes, including nutrient cycling, pollination, pest control and decomposition. The response of invertebrate assembles is of great scientific interest, as studies of post-rodent eradication recovery on islands have typically focused on endangered birds, mammals or reptiles (St Clair et al., 2011). Where previously conducted, post-rodent eradication invertebrate surveys have documented highly heterogeneous impacts, ranging between negligible and substantial increases in abundance, as well as both increases and decreases in diversity (Green, 2002; Samaniego-Herrera et al., 2017; Sinclair et al., 2005; Van Aarde et al., 2004). In several of these studies, the impacts of rodent eradication were somewhat obscured by limited sample sizes and/or temporal variability, emphasising the need for comprehensive, large-scale surveys. LHI is a suitable model system for further research of invertebrate responses to rodent eradication, as invertebrates are exceptionally abundant and diverse, with > 1600 described species, and it is unclear how different taxonomic groups were historically impacted by rodents (Cassis et al., 2003).

Here we present the results of comprehensive invertebrate sampling undertaken on LHI before and after the REP to examine whether: 1) total and Order-level invertebrate abundance increased following the REP; and 2) Order-level assemblage composition changed following the REP.

## 2. Methods

### 2.1. Surveys

Invertebrate sampling was conducted at 20 sites across the lowlands on LHI (Figure 1). To ensure balanced representation of both major soil types on the island – which correspond with a suite of environmental variables such as topography and forest type (Sheringham et al., 2016) – 10 sites were located on calcarenite bedrock and 10 were located on basalt bedrock (Figure ***1***; Supplementary Table S1). Each site spanned an area of *ca.* 100 m^2^, and consisted of various tree, shrub and associated groundcover species, and a substrate with varying amounts of leaf litter, soil and bare rock. Surveys were conducted over two 12-month periods, one before (September 2016 – August 2017) and one after (September 2023 – August 2024) the eradication of rodents. In each period, we undertook collections every three months at the beginning of each Austral season (i.e., in December, March, June and September)

**Figure 1.**
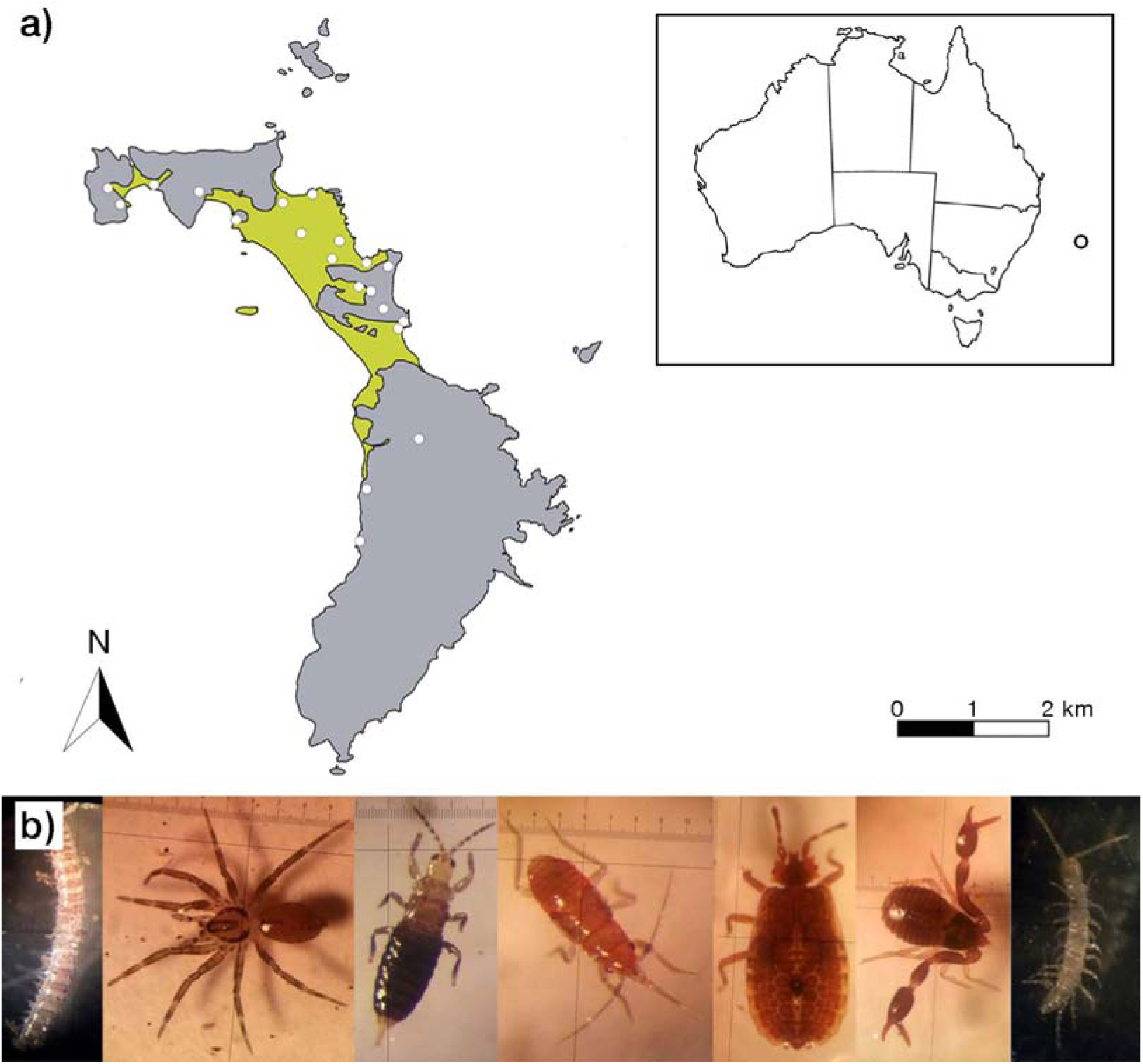
**a)** Locations of invertebrate survey sites on Lord Howe Island. Green shading: calcarenite bedrock, grey shading: basalt bedrock. Inset: location of Lord Howe Island relative to Australia. **b)** Examples of Lord Howe Island-endemic invertebrates. Photographs by Author 5.

At each site, we collected invertebrates from three different microhabitats. First, wraps and blocks were deployed on two adjacent trees with trunk diameter greater than 25 cm at breast height. Tree wraps were designed to mimic exfoliating bark, and consisted of a piece of corrugated cardboard (560 mm × 280 mm) with a piece of waxed cardboard stapled to the outside to increase the longevity of the wrap. Both layers were wrapped around the trunk of each tree (**Error! Reference source not found.**). Two wraps were strapped to each tree on opposite sides of the trunk. Tree blocks were designed to mimic small tree hollows. Each block consisted of a 150 mm wide × 200 mm long × 50 mm deep piece of pine wood with a 95 mm wide keyhole-shaped cavity recessed into the surface facing the tree, with a 25 mm wide opening facing towards the ground (Figure 2). Four blocks were strapped to each tree. In two of these, the cavity was recessed to a depth of *ca.* 17 mm (permitting the entry of mice), and in two the cavity was *ca.* 9 mm deep (preventing access by mice).

**Figure 2.**
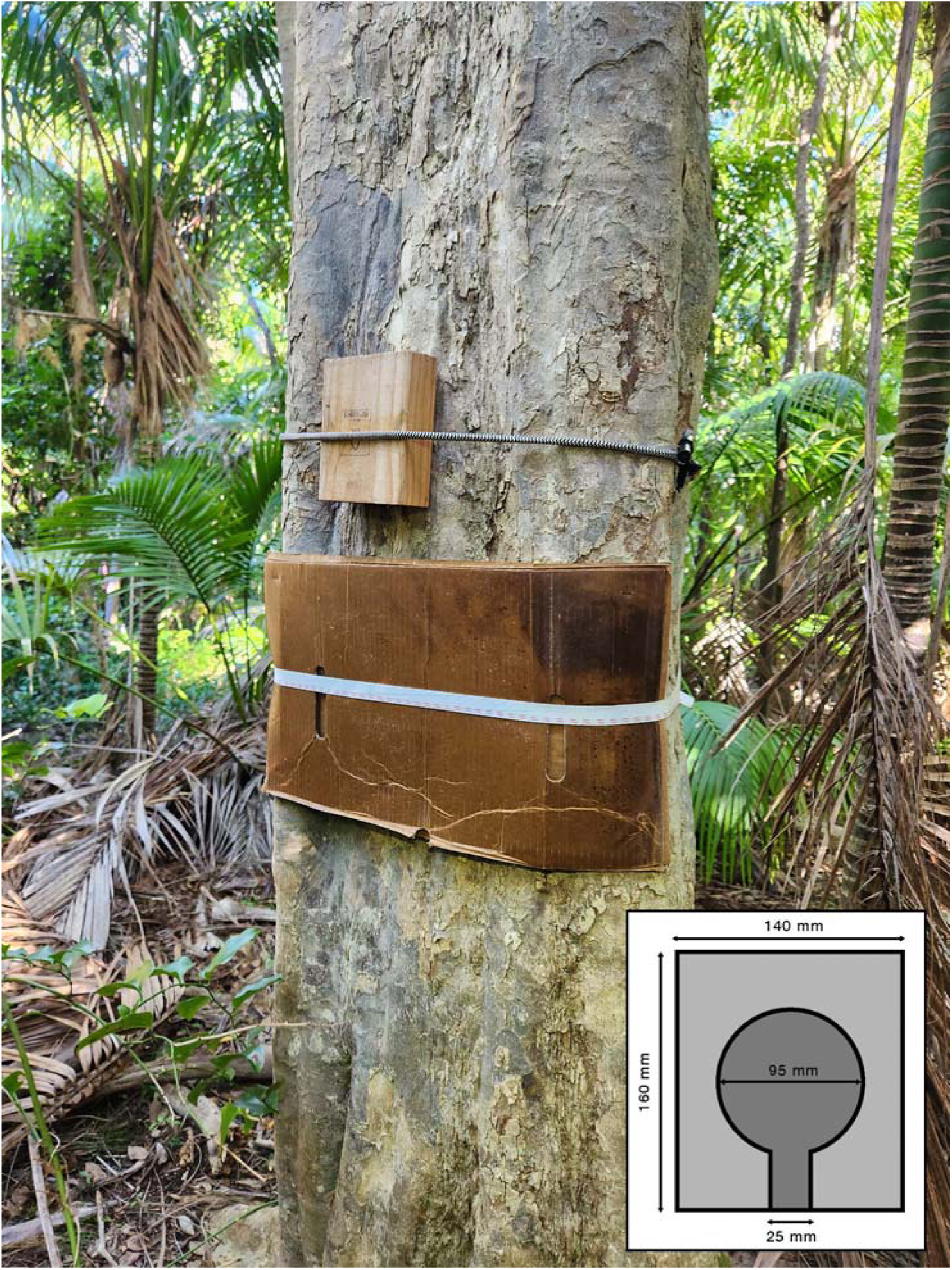
Cardboard tree wrap and wooden tree block strapped to a tree to mimic exfoliating bark and a tree hollow, respectively. Inset: schematic representation of a tree block. Dark shaded region incised to a uniform depth of either 9 mm or 17 mm. When deployed, the incised surface was positioned facing inwards against the tree with the opening directed downwards.

Second, a “roach hotel”, designed to mimic rotting vegetation and used in previous Blattodea studies on LHI (Carlile et al., 2018), was placed directly onto the forest floor between each sampled tree and in direct contact with soil substrate. The roach hotel consisted of three layers of corrugated cardboard (560 mm × 280 mm) topped by a piece of waxed cardboard. The cardboard pieces were held together with strapping tape and dampened with fresh water immediately before deployment. Third, we collected four leaf litter samples from each site: one approximately 3 m to the northern side and one approximately 3 m to the southern side of either tree. Leaf litter samples were collected at the time of recovery of other sampling equipment.

Tree wraps, tree blocks and cockroach hotels were deployed for three months before invertebrate collection. To harvest invertebrates from the tree wraps, a plastic skirt was attached around the tree to catch any dislodged animals. The cardboard wraps were then removed and any invertebrates residing between the cardboard and the trunk were collected either by hand, with forceps, or with the use of a manual invertebrate vacuum. In some cases, larger, fast-moving invertebrates were immobilised with Servisol® electronic freezer spray before capture. The Order and size of any escaping invertebrates were also recorded. Blocks were removed individually, and any residing invertebrates were collected as above. The cockroach hotel was dismantled by sequentially removing individual cardboard layers and retrieving any invertebrates present. Individuals located at the interface between the hotel and the soil were also collected. Finally, leaf litter samples were collected by pressing a stainless-steel circular cutter (25 cm in diameter) to the ground to cut through leaf litter, and retrieving all contained vegetative material and invertebrates into 150 um plastic bags. All other invertebrates captured were immediately placed into Ziploc® bags and wrapped loosely to prevent damage. Bags were sealed and refrigerated prior to processing. After each seasonal round of sampling, the sites were reset as above using a new pair of trees.

### 2.2. Sorting and identification

Samples were manually sorted and identified to Order. To extract invertebrates from leaf litter, the litter was emptied into a Tullgren funnel and placed above a collection jar containing 75% ethanol, then kept under a heat lamp for 48 hr (a sufficient period to dry leaf litter fully). Body measurements were taken using digital callipers or a ruler placed below a petri dish (measured to the nearest millimetre from the anterior margin of the head to the posterior margin of the abdomen). After identification, all samples were preserved in 100% ethanol or frozen at −20 °C. As a result of irregularities in collection, such as clumped distributions or large numbers of escapees, the orders Collembola and Polyxenida, and the family Formicidae, were excluded from statistical analyses. In addition, due to ongoing systematic uncertainty (Van Dam *et al*. 2018) and the subtlety of morphological differences, mites were only identified to the Superorder Acariformes.

### 2.3. Statistical analyses

Analyses were undertaken using *R* v.4.4.2 in RStudio v. 2024.12.0. All invertebrates collected in a season from a given site were pooled across sampling methods (tree wraps, tree blocks, cockroach hotels and leaf litter; each such pool is hereafter referred to as a distinct “collection”). We first examined changes in total invertebrate abundance, diversity and assemblage composition. For each collection, we calculated total invertebrate abundance and Ordinal diversity. Ordinal diversity was estimated using three metrics: richness, the Margalef Index (a standardised measure of richness that accounts for sample size) and the Simpson Index (an estimate of diversity that incorporates the relative abundances of Orders). Unidentified invertebrates were included in our estimates of total abundance; however these were removed from all other analyses. Diversity metrics were calculated in the package *vegan* v.2.6.10 (Oksanen et al., 2013). We used type-III analyses of variance (ANOVAs) in the package *lme4* v.1.1.37 (Bates et al., 2015) to test whether total abundance, richness, Margalef diversity and Simpson diversity varied based on pre- or post-eradication status, Austral season and/or soil type. Sampling site was included in our models as a repeated measure, and we used Satterthwaite’s method in *lmerTest* v.3.1.3 (Kuznetsova et al., 2017) to test significance. All interactions were also tested, but only significant terms are reported. In addition, to determine which of the factors and/or interactions best fit the data, we undertook stepwise model decomposition and selection based on the Akaike Information Criterion in the R package *MASS* v.7.3.65 (Ripley et al., 2013). As this method does not consider repeated measures, site was not included as a factor.

We assessed whether the relative abundances of all the Orders varied pre-versus post-eradication, as well as between seasons, using a two-way permutational multivariate analysis of variance (PERMANOVA) in *vegan*. We conducted post-hoc comparisons of diversity in corresponding seasons before and after rodent eradication using one-way PERMANOVAs with Bonferroni corrections. The similarity of invertebrate assemblages from individual collections was also visually summarised using non-metric multi-dimensional scaling (NMDS) in *vegan*. NMDS was chosen to account for the non-independence and non-linearity of the data (based on preliminary analyses).

Due to the lack of temporal replication of season in our pre- and post-eradication samples, we implemented two methods to test whether our results were confounded by factors other than the rodent eradication itself. First, we checked whether changes in insect abundance may have been driven by climatic conditions by testing for a correlation between abundance and rainfall in the preceding month. Rainfall data were collated from the Bureau of Metrology database (Bureau of Meteorology, 2023). Second, to more finely isolate the impacts of rodent presence, we filtered the data to include only invertebrates with body length > 13 mm (hereafter “large” invertebrates; following Watts et al. 2020), as these are subject to significantly higher levels of rat predation (St Clair et al. 2011). The previously described ANOVAs were re-run using the size-filtered data.

Finally, we examined changes in the eight Orders with the highest total abundances across all years and seasons. Using complete abundance data (i.e. not filtered for body length), we used two-way repeated-measure ANOVAs to test whether the abundance of each Order varied pre- and post-eradication, and/or between seasons.

## 3. Results

### 3.1. Total abundance and Ordinal diversity

We identified a total of 33 Orders across all collections (*n* = 25 before eradication and *n* = 28 after eradication; 36 Orders were present when including Collembola, Lepidoptera and Polyxenida). 24209 invertebrates were collected in total, with 9380 collected prior to eradication and 14829 collected after eradication (Figure 3a). Across all collections, the eight most abundant Orders were (in descending order) Isopoda, Araneae, Blattodea, Coleoptera, Psocodea, the Superorder Acariformes, Polydesmida and Diptera, which together comprised 89% of all captured invertebrates.

**Figure 3.**
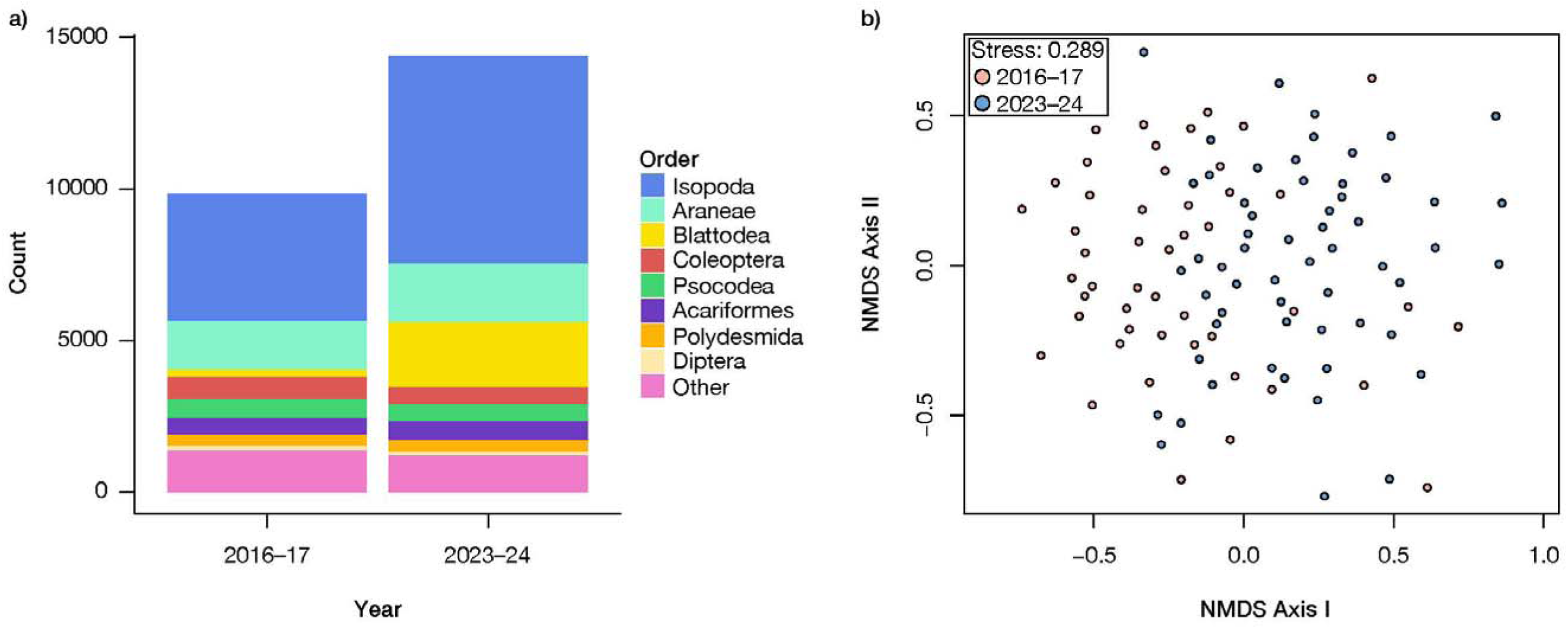
**a)** Proportions of invertebrate Orders before and after rodent eradication. The eight most abundant Orders are labelled. **b)** nMDS plot of invertebrate assemblages from all individual collections. Colours denote year of collection.

Total invertebrate abundance increased significantly after the rodent eradication (F_1,126_ = 8.927, *p* = 0.003; Figure 4a). Abundance also varied significantly between seasons (F_3,144_ = 4.929, *p* = 0.003; Figure 4a).

**Figure 4.**
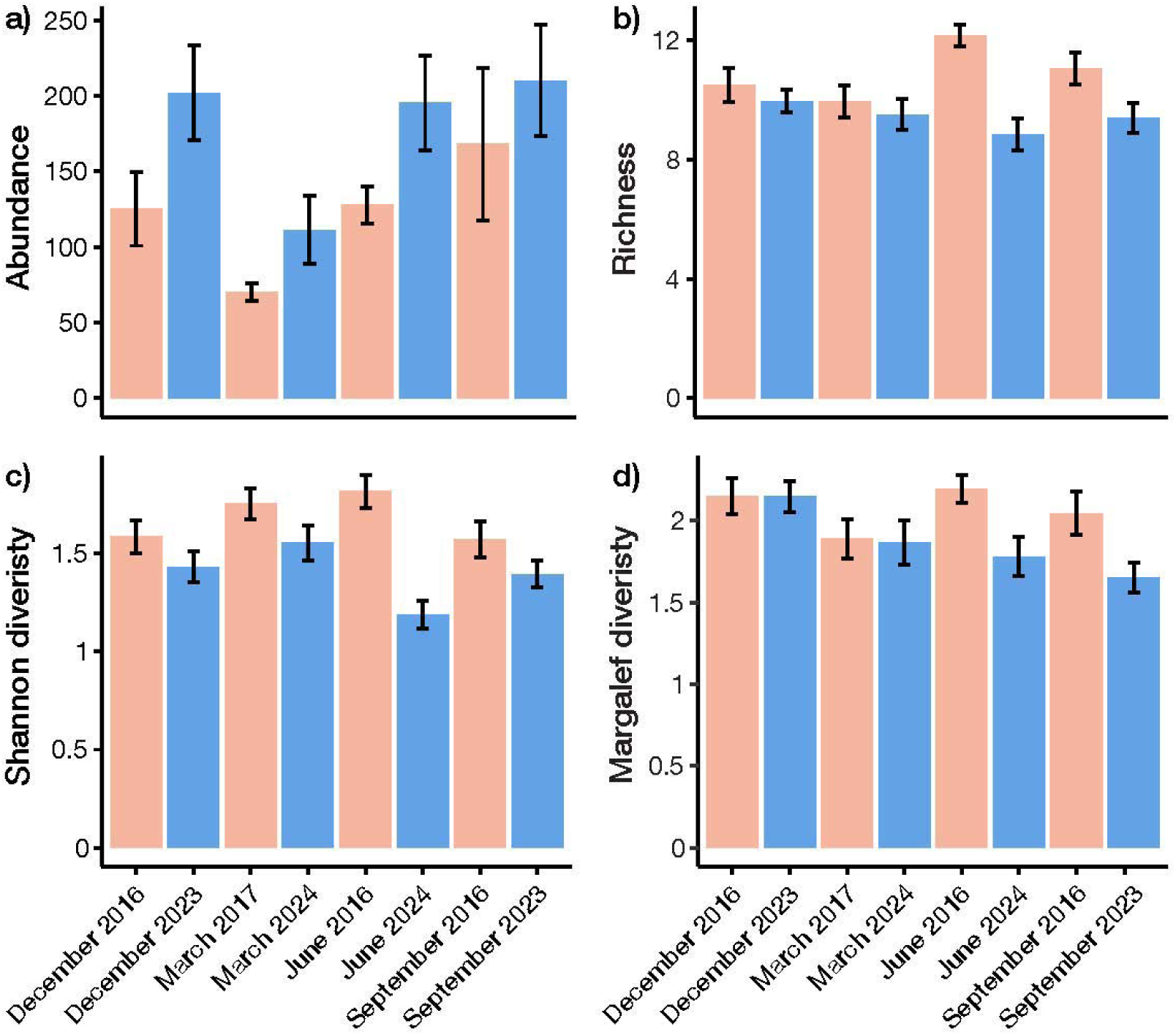
Mean (± S.E.) abundance and Ordinal diversity of invertebrates per site, stratified by season, before and after the eradication of rodents. **a)** Total abundance. **b)** Ordinal richness. **c)** Shannon diversity. **d)** Margalef diversity.

There was a significant interaction between the effects of year and season upon Ordinal richness (F_3,126_ = 4.437, *p* = 0.005; Figure 4b), where richness was lower in Autumn (March) post-eradication but constant across other seasons, as well as between season and soil type (F_3,126_ = 3.345, *p* = 0.020). There was a similar, significant interaction between year and season upon Shannon diversity (F_3,126_ = 4.778, *p* = 0.003; Figure 4c), whereby diversity only varied significantly in Autumn, with a decrease post-eradication. In contrast, Margalef diversity was significantly post-eradication (F_1,126_ = 7.482, *p* = 0.007; Figure 4d), as well as significantly different between seasons (F_3,126_ = 3.200, *p* = 0.026), with highest overall values in Spring. Stepwise model decomposition indicated that variations in abundance, richness, Shannon diversity and Margalef diversity were all best explained by a combination of year and season (with or without soil type), with variable interactions among these factors (Supplementary Table S1).

The relative abundances of all Orders varied across collections with a significant interaction between year and season (PERMANOVA: F_3,152_ = 2.356, *p* < 0.001). Post-hoc analyses revealed significant differences between years in each of the four seasons (Supplementary Table S2). NMDS ordination supported this finding, with site-specific assemblages from 2016–17 and 2023–24 forming two relatively distinct clusters (Figure 3b). No groupings were apparent based on season or soil type.

There was no significant relationship between total invertebrate abundance and rainfall in the preceding month, albeit based on a limited sample size of 8 data points (*t* = 2.249, *p* = 0.066, *r^2^* = 0.367; Supplementary Figure S1). When the data were filtered for body lengths > 13 mm, there was a significant increase in the abundance of large invertebrates after the rodent eradication (F_1,126_ = 19.791, *p* < 0.001; Figure 5a), with a more pronounced increase than was evident when small and large invertebrates were analyses together. No other factors impacted abundance. Ordinal richness of large invertebrates also increased significantly following the eradication (F_1,126_ = 10.948, *p* = 0.001; Figure 5b), unlike the non-size-filtered data, where Ordinal richness did not clearly shift after the eradication. Shannon diversity of large invertebrates varied significantly between seasons (respectively: F_3,126_ = 3.015, *p* = 0.033), with greatest values in Winter (June), but did not change significantly pre-versus post-eradication. Margalef diversity of large invertebrates varied according to significant interaction between year and soil type (F_1,126_ = 5.892, *p* = 0.017), but without any clear main effects of the eradication or season.

**Figure 5.**
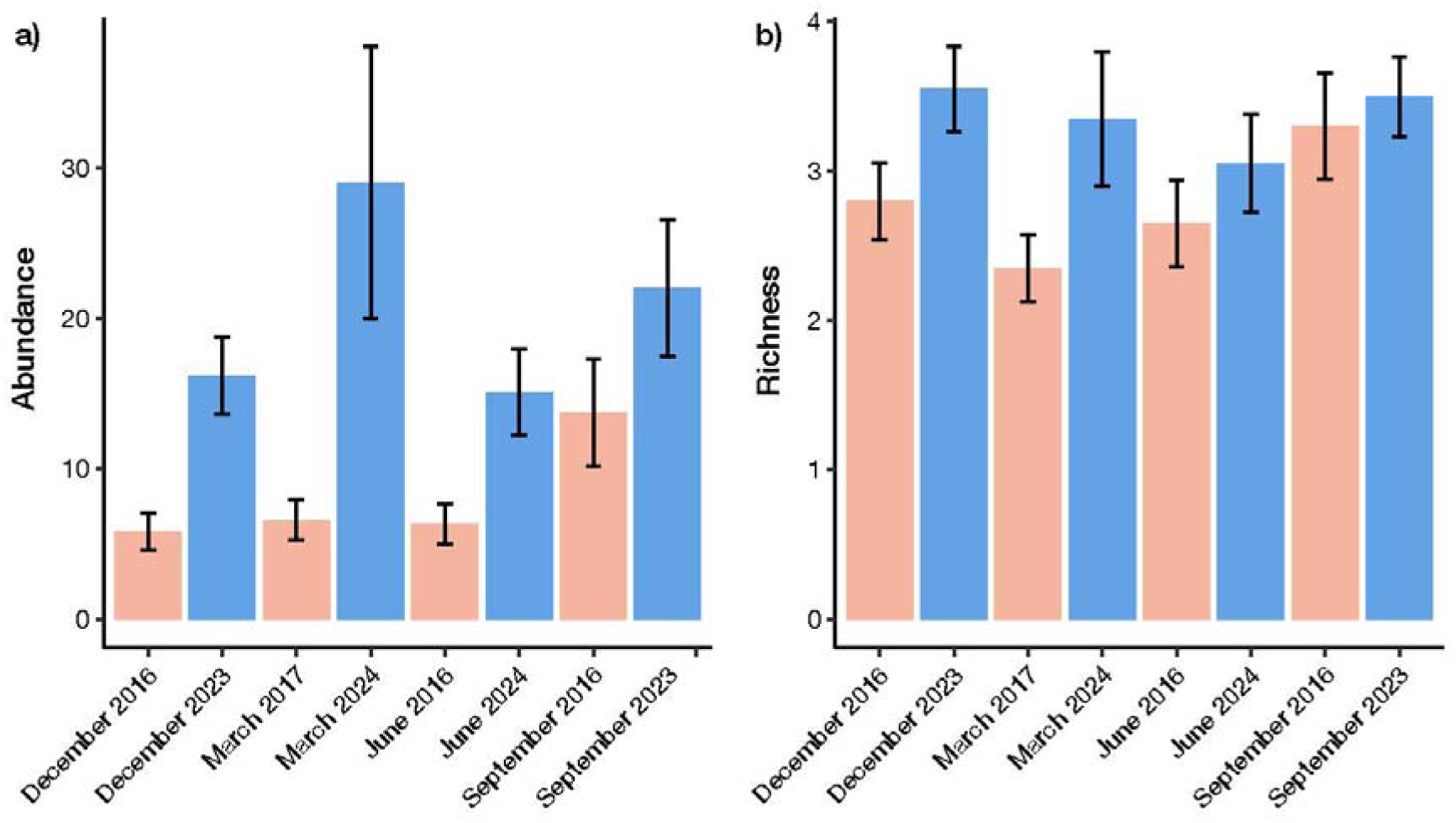
Mean (± S.E.) abundance and diversity of large invertebrates per site, before and after the eradication of rodents in each season. **a)** Abundance. **b)** Ordinal richness.

### 3.2. Individual Orders

The abundance of Isopoda increased significantly following rodent eradication (F_1,133_ = 4.399, *p* = 0.038; Figure 6a), with no significant effect of season (F_3,133_ = 2.652, *p* = 0.051). The abundance of Araneae varied based on a significant interaction between year and season (F_1,133_ = 4.399, *p* = 0.038; Figure 6b), whereby abundance increased following the eradication in Winter only. The abundance of Blattodea increased significantly following eradication (F_1,133_ = 21.187, *p* < 0.001; Figure 6c), without any significant variation between seasons (F_3,133_ = 1.855, *p* = 0.194). There was no significant change in Coleoptera numbers before versus after rodent eradication (F_1,133_ = 3.171, *p* = 0.077; Figure 6d), nor between seasons (F_3,133_ = 0.594, *p* = 0.620). There was a significant interaction between year and season upon the abundance of Psocodea (F_3,133_ = 4.906, *p* = 0.003; Figure 6e), with a substantial increase in Spring following rodent eradication but a decrease in Autumn. For Acariformes, there was insufficient variance based on site, thus we proceeded with a standard (unnested) linear model. We found that Acarina numbers varied according to a significant interaction between year and season (F_3,152_ = 9.4411, *p* < 0.001; Figure 6f), whereby post-eradication abundance was significantly lower in Autumn, but significantly higher in Spring (September). The abundance of Polydesmida did not vary pre-versus post-rodent eradication (F_1,133_ = 0.040, *p* = 0.843; Figure 6g) or between seasons (F_3,133_ = 1.945, *p* = 0.314). Finally, we also modelled the abundance of Diptera with a standard linear model, which also found no significant changes in abundance following rodent eradication (F_1,152_ = 0.538, *p* = 0.465; Figure 6h) or between seasons (F_3,152_ = 1.507, *p* = 0.215).

**Figure 6.**
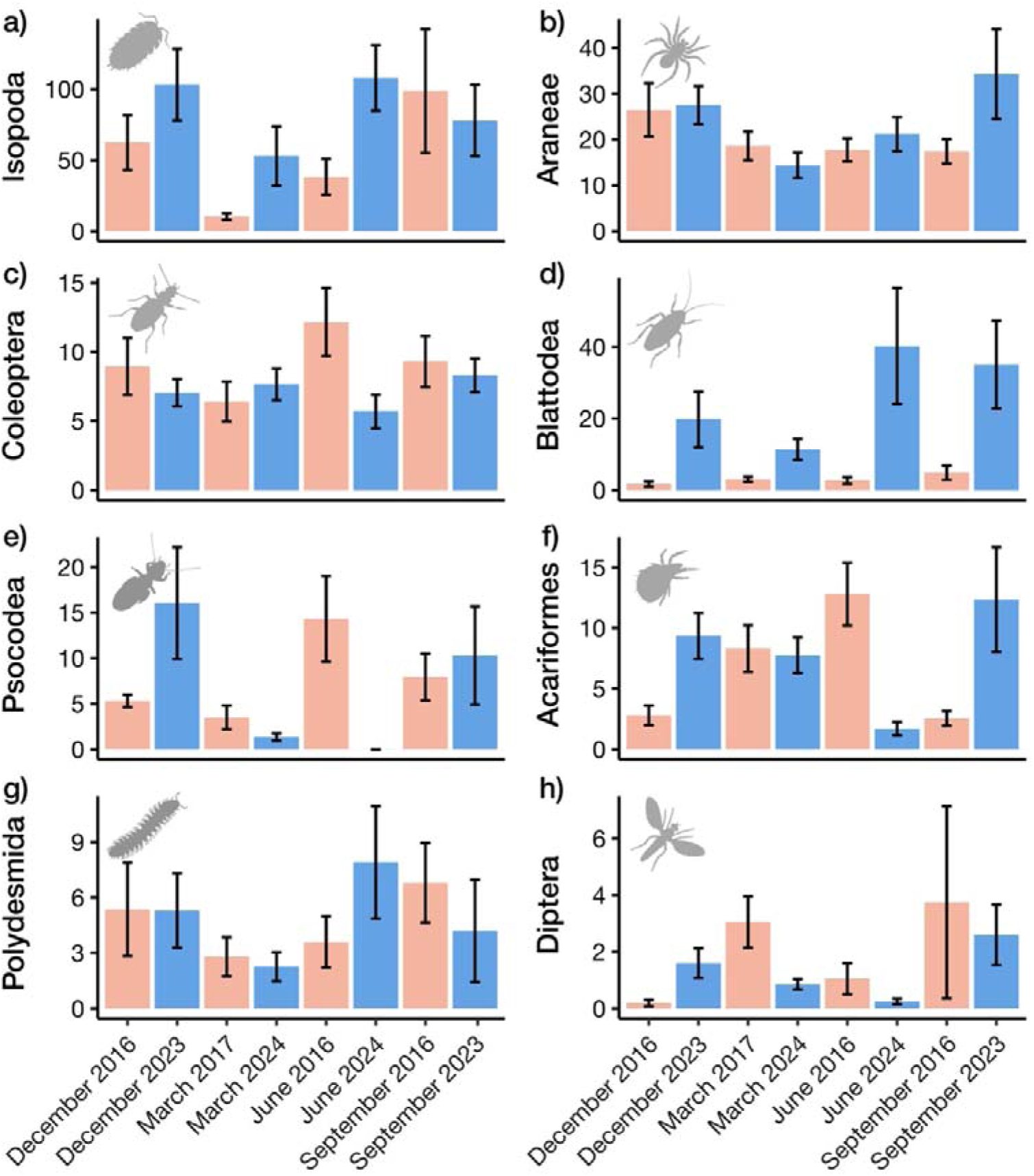
Mean (± S.E.) abundance of specific Orders per site, before and after the eradication of rodents in each season. Inset graphics sourced from PhyloPic.

## 4. Discussion

This study augments the few previous investigations of island invertebrate responses to rodent eradication (Green, 2002; Samaniego-Herrera et al., 2017; Sinclair et al., 2005; Van Aarde et al., 2004). Overall, we found clear evidence of increased total abundance, and changes in assemblage composition, following rodent eradication. This reinforces the broad and substantial biodiversity benefits of invasive predator eradication, and also demonstrates a meaningful return on investment from the eradication. We acknowledge that the lack of temporal replication before and after eradication introduces potential confounds on the results, and limits our ability to definitively link ecosystem shifts to rodent eradication alone. However, the weak and non-significant correlation between rainfall and abundance suggests that the impact of this important climatic variable was slight, although we cannot exclude an impact of other environmental changes in the years between our samples. Furthermore, by examining only the species that are probable targets for rodent predation (based on body size > 13 mm; St Clair, 2011), we found larger and more consistent increases in abundance following the eradication, as well as modest – but significant – increases in Ordinal richness. Together, these results suggest that the broad increase in abundance was at least partially driven by the eradication of rodents, and that the rodent-suppressed component expanded especially substantially. That the abundance and diversity of insects were also strongly impacted by seasonal variation is consistent with some previous findings (Sinclair et al., 2005) and highlights the non-linearity of ecosystem recovery.

Due to the long history of rodent presence on LHI, it has remained an open question how current invertebrate assemblages differ from those existing before rodents arrived. Previously known impacts have been primarily at the species level, and include the loss of two endemic land snails (*Epiglypta howinsulae* and *Placostylus bivaricosus etheridgei*; Ponder 1997), 11 beetle species (C. Reid, unpub. data) and the LHI population of the phasmid *Dryococelus australis* (now restricted to Ball’s Pyramid; Priddel et al., 2003). Our comparisons of dominant Orders (which represent *ca.* 90% of total abundance) revealed that Blattodea and Isopoda exhibited the largest increases following the REP, while the abundance of Coleoptera and Polydesmida did not change. Interestingly, Coleoptera have the most documented examples of post-rodent rebound on islands of any invertebrate Order (reviewed by St Clair, 2011), although this result may be an artefact of the group’s high species richness. The preferred prey of invasive rodents vary greatly between islands and often entail complex indirect effects (Harper & Bunbury, 2015; St Clair, 2011). Dietary preferences of black rats and house mice on LHI have not been studied, hence it is challenging to explain why some Orders were more or less impacted on LHI.

Despite the shifts in abundance and assemblage structure, there were no clear changes in total Ordinal diversity, irrespective of the metric used. It appears that reciprocal increases and decreases across Orders maintained similar levels of richness and evenness between years. Interestingly, we did find a significant increase in Ordinal richness among large invertebrates, suggesting that these shifts may have favoured larger taxa in particular. While our coarse taxonomic resolution precludes comment on changes at the species level, even more granular studies of invertebrate communities have not consistently found increases in diversity following rodent eradications, suggesting that the present results may not be an artefact of taxonomic resolution (e.g. Green, 2002; Samaniego-Herrera et al., 2017; Sinclair et al., 2005; Van Aarde et al., 2004; but see Palmer & Pons, 1996). Small or non-existent increases in prey diversity are not unusual following invasive predator removal on islands, as population growth may be suppressed by interspecific interactions, resource limitation or rebounds of native predators (Baker et al., 2020; Bird et al., 2024; Borrelle et al., 2018; Kurle et al., 2008; Towns et al., 1997). In concordance, the abundance of native predators appears to have risen following rodent eradication on LHI, as our opportunistic captures of geckos revealed a two-fold increase in their abundance (unpub. data), and similar increases in insectivorous birds have also been noted elsewhere (O’Dwyer et al., 2023).

This study offers a broad view of invertebrate recovery on LHI. The overall pattern points towards a turnover in invertebrate assemblage compositions, as has been observed following rodent eradications on Kapiti Island (Sinclair et al., 2005), Mexican tropical islands (Samaniego-Herrera et al., 2017) and Macquarie Island (M. Houghton, pers. comm.). The long-term ecological consequences of such shifts are difficult to predict. While ecological niches and functional groups were not recorded, the Orders with the largest increases consist primarily of detritivores and decomposers, which may have downstream implications for nutrient cycling. In addition, we cannot exclude the possibility that invasive invertebrate species are among those experiencing significant increases, as predator eradications often do not restore ecosystems to their pre-disturbance trajectories (e.g., Bode et al., 2015). On nearby Blackburn Island, habitat restoration has inadvertently allowed for the establishment of invasive isopods (Author 2, pers. obs.). Future sampling, replicating the present design, would be invaluable for monitoring the development of the invertebrate assemblages, and would also provide more robust temporal replication, which would help to affirm or temper the conclusions outlined here. A planned genetic study using the samples collected here will provide additional detail on the invertebrate diversity now present on LHI.

Our invertebrate sampling was a single component of a broader biodiversity monitoring program. It was previously found that the rodent eradication allowed for the expansion of formerly bottlenecked populations of the cockroach *Panesthia lata* (Adams et al., 2024) and had a major positive impact on the breeding success of black-winged petrels (*Pterodroma nigripennis*; O’Dwyer et al., 2023). The removal of rodents also benefitted a range of land birds, including Lord Howe woodhens which have increased more than seven-fold since the REP (O’Dwyer et al., 2024). This species primarily consumes soil dwelling invertebrates (Miller & Mullette, 1985) and their rapid increase following the eradication may have been facilitated by the increase in large invertebrates. Conversely, the rapid increase in woodhen abundance might be limiting the growth of some invertebrate populations. The contrast between the heterogeneous shifts in invertebrate assemblages and the more straightforward increases in bird abundance is notable but not unprecedented (reviewed by St Clair, 2011; Watari et al., 2011). Birds were historically top predators on LHI and their recovery is presumably less impacted by mesopredator release. It is nonetheless promising that changes are already evident across trophic levels, as the benefits of rodent eradication can require up to decades to manifest, especially for indirectly impacted taxa (Bird et al., 2024; Jones, 2010; Philippe□Lesaffre et al., 2023; Towns, 2009). Further planned components of the biodiversity benefits program include an assessment of little shearwater (*Puffinus assimilis*) breeding success and a focused study of the impacts on geckos, which will provide valuable insights as LHI continues to recover.

Invertebrates are often omitted from environmental monitoring programs, including on islands (reviewed by Samaniego-Herrera et al., 2017; Ward & Larivière, 2004). Many groups are poorly described or taxonomically unstable; and even modest collection efforts can yield large sample sizes that require significant effort to process. There is also a lack of standardised collection protocols and balanced sampling regimes (reviewed by Brown & Matthews, 2016; Neville & Yen, 2007; Webb et al., 2022). Despite these challenges, we demonstrate that a relatively simple sampling design, with coarse taxonomic resolution, can illuminate ecosystem dynamics following rodent eradication. This highlights the benefits of the REP as both a conservation and biodiversity management measure, as well as an experiment to examine the consequences of rodent invasion.

## Acknowledgements

We thank the Lord Howe Island Board, and staff members including Anthony Wilson and Shane Deaco, for assistance with collection. We also acknowledge the Australia and Pacific Science Foundation and the Australian Research Council for support of our research on Lord Howe Island. The raw data and R code used to prepare this manuscript are to be made available on Dryad.

