## Supplementary Material for "Increases in invertebrate abundance and shifts in assemblage composition following Rodent Eradication on Lord Howe Island"

**Results**

**Supplementary Table S1.** Results of stepwise model decomposition analyses. The best-fitting model was found based on the Akaike Information Criterion in *MASS*.

| **Variable** | **Best-fitting model** |
| --- | --- |
| Total abundance | Year + soil + season + soil * season |
| Ordinal richness | Year + season + soil + year * season + season * soil |
| Shannon diversity | Year + season + year * season |
| Margalef diversity | Year + season + soil + year * season + year * soil |

**Supplementary Table S2.** PERMANOVA results comparing the overall abundances and relative frequencies of invertebrate Orders between 2016–17 and 2023–24, per season. Asterisk (*) denotes a statistically significant *p*-value.

| **Season** | **F-statistic** | **Degrees of freedom** | ***p*-value** |
| --- | --- | --- | --- |
| Spring | 3.519 | 1, 38 | 0.008 * |
| Summer | 3.293 | 1, 38 | 0.003 * |
| Autumn | 7.705 | 1, 38 | < 0.001 * |
| Winter | 2.664 | 1, 38 | 0.015 * |

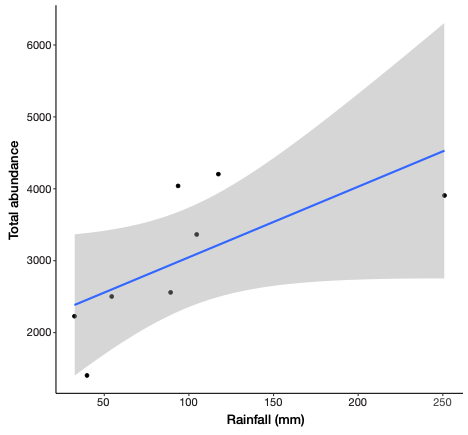

**Supplementary Figure S1.** Relationship between rainfall in the month preceding collection and total invertebrate abundance summed across all 20 sites. Blue line represents line of best fit, grey band indicates 95% confidence interval.
